# Signatures of Selection in Composite *Vrindavani* Cattle of India

**DOI:** 10.1101/2020.10.19.344747

**Authors:** Akansha Singh, Arnav Mehrotra, Cedric Gondro, Andrea Renata da Silva Romero, Ashwni Kumar Pandey, A Karthikeyan, Aamir Bashir, B.P. Mishra, Triveni Dutt, Amit Kumar

## Abstract

*Vrindavani* is an Indian composite cattle breed developed by crossbreeding taurine dairy breeds with native indicine cattle in the 1960s. About 190,000 semen doses of *Vrindavani* bulls have been distributed to the farmers till date. The animals are under artificial and natural selection for higher milk production and adaptation to the tropical climate, respectively. However, the selection response for production and adaptation traits in the *Vrindavani* genome is not explored. In this study, we provide the first overview of the selection signatures in the *Vrindavani* genome. 96 *Vrindavani* cattle were genotyped using the BovineSNP50 BeadChip and the SNP genotype data of its constituent breeds were collected from a public database. Within-breed selection signatures in Vrindavani were investigated using the integrated haplotype score (iHS). Vrindavani was also compared to each of its parental breeds to discover between-population signatures of selection using two approaches, cross-population extended haplotype homozygosity (XP-EHH) and fixation index (F_ST_). Selection of signature identifies 11 common region identified by more than one harbouring genes such as *LRP1B*, *TNNI3K*, *APOB*, *CACNA2D1*, *FAM110B and SPATA17* associated with production and adaptation. Overall, our results suggested stronger selective pressure on regions responsible for adaptation compared to milk yield.

## 1. Introduction

The benefits of crossbreeding between high yielding *Bos taurus* and environmentally resistant *Bos indicus* breeds in tropical production systems have been well established over the last half-century. Crossbred cattle have played an important role in meeting India’s rising demand for milk. Despite constituting only 20.7% of India’s milch herd, the crossbreds contribute 26% of India’s 187.75 MT annual milk production (Livestock Census, 2020; DAHD&F, 2018-19).

Efforts to determine the optimum levels of *Bos Taurus* inheritance in different geographical tracts of India started in the 1950s (Amble and Jain, 1967), with varying levels of success. Among these was a four breed crossing scheme initiated at the Indian Veterinary Research Institute in 1968. Briefly, a foundation stock of 400 indigenous Hariana cattle was inseminated with Holstein Friesian (HF), Jersey and Brown Swiss (BSW) semen to produce three genetic groups *viz*., 1/2 Hariana × 1/2 HF, 1/4 Hariana × 1/2 HF × 1/4 BSW, and 1/4 Hariana × 1/2 HF × 1/4 Jersey. These genetic groups were evaluated for production, reproduction and envrionmental adaptation for seven generations. This was followed by inter-mating and selection to create the present day sythetic breed *Vrindavani,* having 25-50% *Bos indicus* and 50-75% *Bos taurus* inheritance (Singh et al., 2011).

For a breeding program to be successful, it is important to constantly evaluate the consequences of the ongoing natural and artificial selection. Over the last decade, SNP microarrays and whole genome sequencing technology has enabled researchers to explore the genetic architecture and signatures of post-admixture selection in synthetic breeds (Cheruiyot et al., 2018; Decker et al., 2014; Kim and Rothschild, 2014). In *Vrindavani* cattle, the Bovine SNP50K array has recently been used to investigate the population structure of the breed (Ahmad et al., 2020; Chhotaray et al., 2019).

In this study we used two complimentary approaches to detect the genomic regions under selection in *Vriandavani* cattle. First, the integrated haplotype score (iHS) was used to detect within-population selection signatures. Second, we compared *Vrindavani* to Hariana, HF, Jersey and BSW by haplotype based (XP-EHH) and single SNP based (*F*_ST_) methods to discover the genomic regions where the synthetic breed has diverged from each of its parental populations after admixture.

## 2. Materials and methods

### 2.1 Sample collection, genotyping and quality conrol

Blood samples were collected from 96 *Vrindavani* cattle kept at the AICRP farm of the ICAR-Indian Veterinary Research Institute, Bareilly, UP (28.3670° N, 79.4304° E), following approval by the Institutional Animal Ethics Committee (IAEC). The Vrindavani bulls are selected and culled on the basis of dam and daughter’s milk yield, respectively. Genomic DNA was extracted and genotyped with the BovineSNP50 v3 BeadChip (AgriGenome Labs Pvt. Ltd., India). Genotypes were called and processed using the GenomeStudio software package (Illumina, Inc.). The SNP coordinates followed the ARS-UCD1.2 assembly of the bovine genome.

To investigate the selection signatures between populations, the genotypic data of Vrindavani’s parent taurine breeds (BSW, HF, Jersey) were accessed using WIDDE (http://widde.toulouse.inra.fr/widde/widde/main.do?module=cattle) and Hariana cattle through KRISHI (https://krishi.icar.gov.in/jspui/handle/123456789/31167) web portal. These included 50K SNP data of HF (**n**◻=◻30), Jersey (**n**◻=◻21), BSW (**n**◻=◻24), and the HD (777K) genotypes of Hariana (**n**◻=◻18). The common SNPs between 50K and HD chip data of Hariana were extracted for further analysis. Due to the large difference in sample sizes, 25 Vrindavani animals out of 96 were retained for the across-population studies. Genotypic data of all the breeds were merged and quality control was performed by filtering non-autosomal and unmapped SNPs. SNPs with less than a 90% call rate, minor allele frequency lower than 0.01 and a significant (P < 0.00001) deviation from Hardy-Weinberg equilibrium were also removed using PLINK (Purcell et al., 2007) leaving 34,197 variants for downstream analysis. The genotypes were phased using BEAGLE v5.1(Browning et al., 2018).

### 2.2 Population structure analysis

The expected heterozygosity (He), observed heterozygosity (Ho) and minor allele frequency (MAF) was estimated using PLINK 1.07 (Purcell et al., 2007). Population relatedness was evaluated using pairwise estimates of *F*_ST_ (Weir & Cockerham, 1984).

Principal Component (PCA) and Admixture (Alexander et al., 2009) analyses were performed to validate the breed separation in our merged dataset. The results were visualised in R with the basic plot function (R Core Team, 2018).

### 2.3. Selection signature analyses

The within population signatures of selection in Vrindavani (**n**=96) were computed using the integrated haplotype score (iHS) (Voight et al., 2006). The ancestral allele information for the iHS test was obtained from Rocha et al. (2014) for the 50k SNP data. The iHS was calculated for each autosomal SNP in Vrindavani through the package rehh (Gautier et al., 2017). Candidate regions were identified using a scan window of 100 kb with a 50 kb overlap. Windows with an average iHS score of 3 (three standard deviations above the mean) or above were considered as candidate regions for selection.

To ascertain between-population selection signatures, Vrindavani was compared to each of its parental populations using XP-EHH and *F*_ST_.

The XP-EHH (Sabeti et al., 2007) scores were calculated for each pairwise comparison using the package rehh, taking the paretnal breeds as the reference population. To detect positive selection in Vrindavani, average XP-EHH scores were computed for 100-kb regions with a 50 kb overlap. Regions with absolute XP-EHH scores of 3 (Three SD above the mean) or above were considered as putative candidate regions.

The pairwise F_*ST*_ was calculated with VCFTOOLS (Danecek et al., 2011), with a sliding window of 100 Kb and a 50 Kb step size. Windows belonging to the top 0.1% of the *F*_ST_ values were considered as potential regions under selection.

Genes in the candidate regions were annotated using the Ensembl Biomart genes database (release 100). Functional and pathway enrichment analysis was performed using DAVID (Huang da et al., 2009). Each positively selected region was cross referenced with the literature.

## 3. Results

### 3.1. Descriptive statistics and population structure analyses

The heterozygosity and MAF values of *Vrindavani* (*Ho*=0.34, MAF=28) were found similar to the European breeds, particularly to HF (Table S1). The genetic relationship between the *Vrindavani* population and its parent breeds was visualised using PCA. The first and second principal components explained 62.3% and 11.7% of the total variation, placing the *Vrindavani* cattle in between the taurine and indicine breeds which is in agreement with their known lineage (Figure S1). They are however noticeably closer to the Holstein cluster than any of the other parental breeds. In concordance with Ahmad et al. (2020), Admixture analysis with K=4 showed that the average breed composition proportions for our population of *Vrindavani* was 42.5%, 26.0%, 17.1% and 14.4% of HF, Hariana, Jersey and BSW, respectively. (Figure S2).

### 3.2. Within population selection signatures in Vrindavani (iHS)

Considering the recent selection history in *Vrindavani* breed, the genomic sweeps were identified using integrated haplotype score (iHS) approach. A total of 46 significant SNPs (iHS ≥3) distributed across 12 autosomes were identified within the candidate regions (Table S2). The clusters of significant SNPs were found on BTA 3, BTA11, BTA14, and BTA15, with the strongest signal on BTA14 (30.35 Mb - 30.44 Mb; iHS =3.9) (Figure 1). The top 10 regions with their iHS values and genes are shown in Table 1. Functional annotation of the selected regions identified candidate genes related with milk production (*APOB*, *ANO3*, *DNMT3A* and *POMC*) and environmental adaptations or immunity (*DNAJC5B* and *FYB2*).

**Figure 1.**
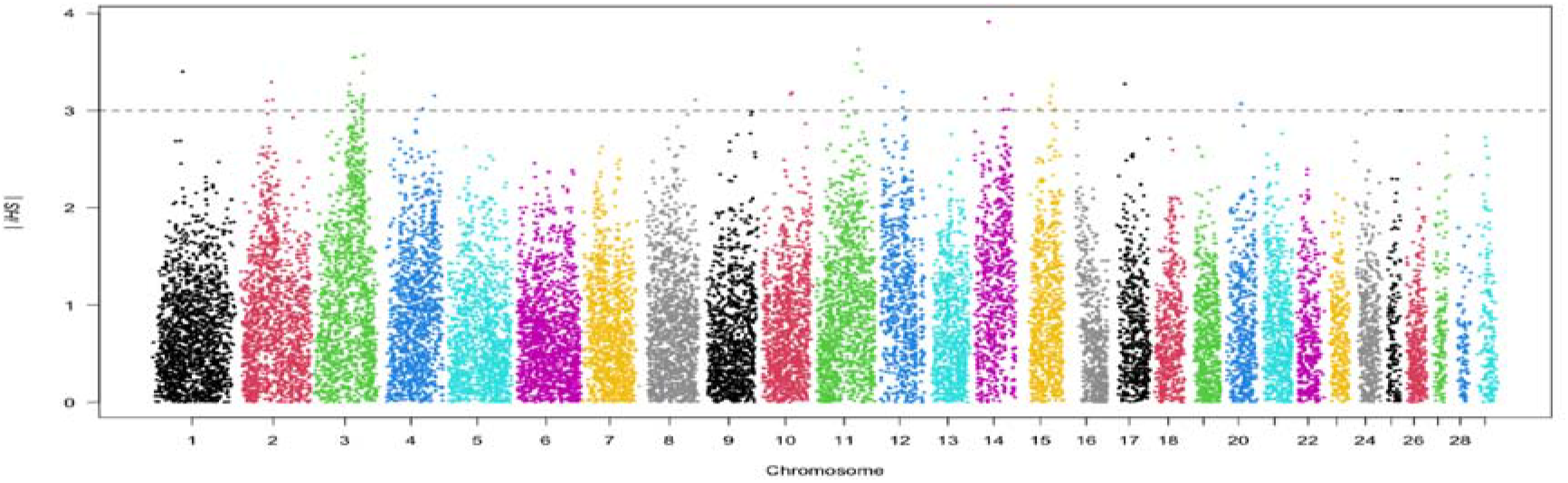
Plot of the integrated haplotype score (iHS) analysis on the *Vrindavani* cattle. The dotted line indicates mean and 3standard deviation as threshold.

**Table 1:** List of the top 10 integrated haplotype score measures (iHS) regions and their closest genes

| BTA | START | END | iHS | GENE |
| --- | --- | --- | --- | --- |
| 14 | 30355171 | 30448182 | 3.913994 | DNAJC5B |
| 11 | 77798022 | 77927967 | 3.631122 | TDRD15, APOB |
| 3 | 89219877 | 89464197 | 3.570568 | C8A, FYB2 |
| 3 | 75039363 | 75644478 | 3.545228 | LRRC7 |
| 11 | 73971560 | 74224519 | 3.483963 | DNMT3A, POMC, EFR3B |
| 11 | 82677576 | 83207199 | 3.404882 | DDX1, NBAS |
| 1 | 57570680 | 57681112 | 3.398117 | CD200R1L |
| 3 | 88677121 | 89136086 | 3.383668 | DAB1 |
| 2 | 61535026 | 61675889 | 3.291714 | LCT, UBXN4, R3HDM1, MIR128-1 |
| 3 | 65667175 | 65807163 | 3.270202 | ADGRL4 |

### 3.3. Across population selection signatures (F_ST_ and XP-EHH)

The Manhattan plot of pairwise XP-EHH analysis between *Vrindavani* and its parent breeds are shown in Figure 2. The selection signals (positive value of XP-EHH for *Vrindavani*), against all parent breeds were detected on 8 autosomes, of which cluster of SNPs are observed on BTA1, BTA2, BTA3, BTA4, BTA11, BTA15, BTA16 and BTA25 (Table S3). BTA4 and BTA3, exhibit the highest number of selected regions in all the breed comparisons. The selective sweep located on BTA4 (91.8 Mb – 92.2 Mb) was detected in comparisons with both BSW and Hariana. It contains the genes *SND1* and *LRRC4* associated with somatic cell count, milk yield and residual feed intake. Two regions on BTA2 (56.4 Mb-56.6 Mb) and BTA25 (31.15 Mb – 31.3 Mb) were detected in comparisons with Hariana and Jersey. It contains *LRP1B* gene and QTLs for somatic cell count and reproduction traits (Cole et al., 2011)

**Figure 2.**
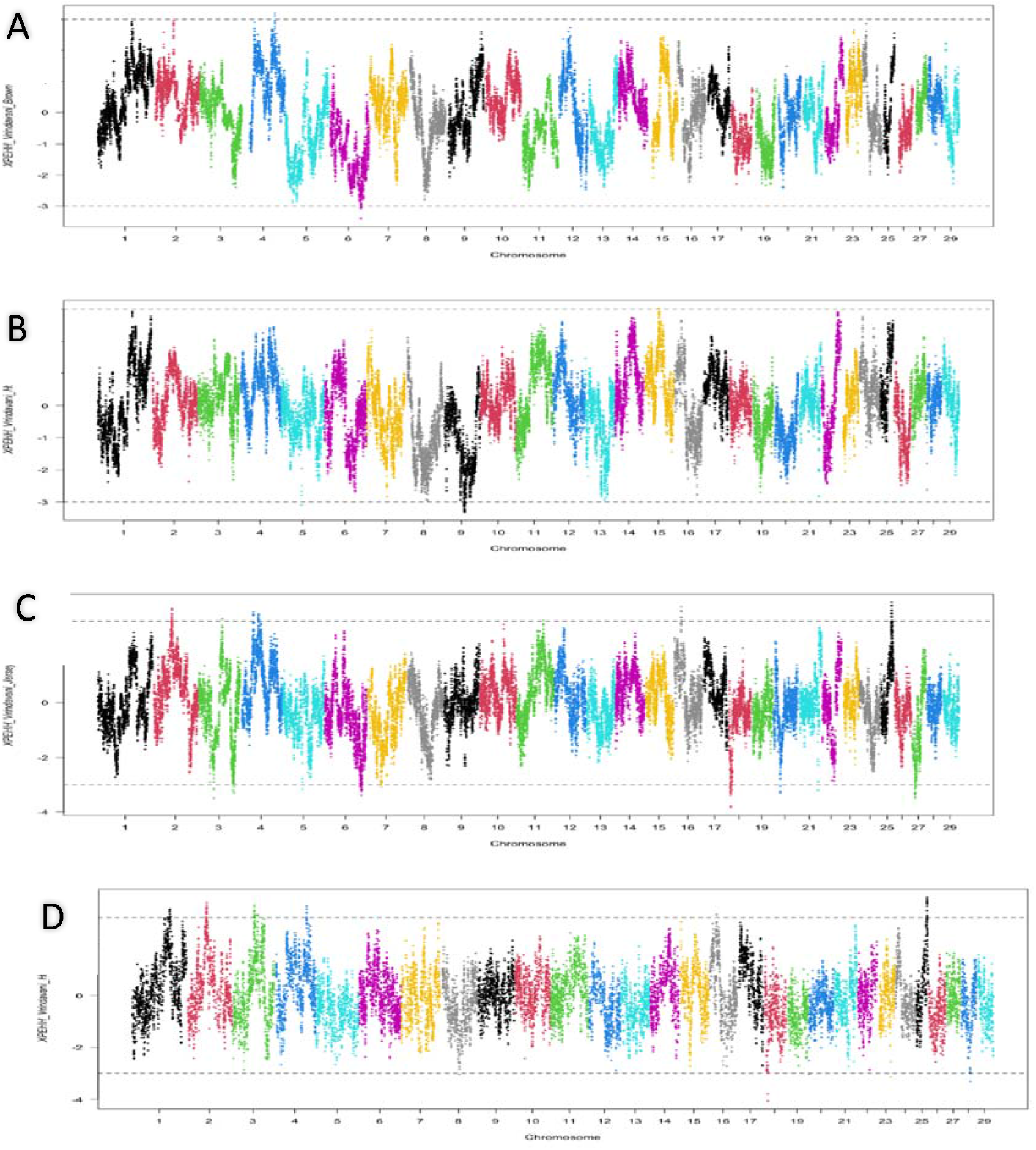
Cross-population extended haplotype homozygosity (XP-EHH) analyses for (A)Brown Swiss, (B) Holstein Friesian (C) Jersey (D) Hariana against the Vrindavani. The dotted line indicates mean and 3standard deviation as threshold.

The mean F_ST_ values of *Vrindavani* in comparison to HF, Jersey, BSW and Hariana were 0.081, 0.110, 0.122 and 0.175 respectively. The pairwise F_ST_ across genome of the *Vrindavani* against its parental breeds were plotted in Figure S3. A total of 124 regions were identified, which were distributed across all autosomes except BTA25 and BTA26 in breed comparisons (Table S4). The regions on the chromosomes having highest F_ST_ values against HF, BSW, Jersey and Hariana were located on BTA4, BTA7, BTA10 and BTA3, respectively.

The F_ST_ signals on overlapping regions located on BTA7 (45.3-48.9 Mb), BTA10 (37.4 Mb - 37.7 Mb) and on BTA14 (12.3 Mb - 12.6 Mb) were observed in comparison with BSW and Jersey, harbouring genes involved with production, reproduction and functional traits (*H2AFY*, *SPOCK1*, *PLA2G4D*, *PLA2G4F* and *GANC*). Two selected regions on BTA14 from 12.3 Mb - 12.6 Mb (against BSW and HF) and 27.3 Mb-27.9 Mb (against Jersey and HF) were identified. These regions include *SPIDR* gene (Scaffold protein involved in DNA repair) associated with milk and milk protein yield) and *NKAIN3* gene (Na+/K+ transporting ATPase interacting 3) related with insulin-like growth factor 1 level.

### 3.4. Comparative analysis of selection signatures

A total of 13 regions on BTA2, BTA3, BTA4, BTA10, BTA11, BTA14, BTA15 and BTA16 were determined by more than one approach; with a region on BTA3 (70.2Mb - 72.2Mb) common to all three approaches (Table 2). Out of 6 regions detected by both the between population approaches (pairwise F_ST_ and XP-EHH), 4 regions were detected in comparisons with taurine breeds (BSW, Jersey and HF); 1 region in comparison with the indicine breed (Hariana) and 1 region against both taurine and indicine breeds(Table2). Functional annotation of the commonly detected regions shows several candidate genes already reported as selection signals or associated with economic traits in different cattle breeds. Genes present in these regions enrich biological processes such as response to virus, ion transportation and post embryonic development, and molecular functions such as ATP binding and motor binding.

**Table 2.** Selection sweeps identified by more than one test in the *Vrindavani* chromosomes (BTA) and annotated genes in these regions

| Test | BTA(Start-End) | Annotated Genes |
| --- | --- | --- |
| XPEHH(Haryana),Fst(Haiana); iHS | 3(70.2-72.18) | TNNI3K |
| XPEHH(Jersey, Haryana);FST(HF) | 2(55.5-56.7) | LRP1B |
| XPEHH(Jersey); Fst(HF) | 4(38.1-40.1) | CACNA2D1, 7SK,SEMA3C |
| XPEHH(Jersey); Fst(HF) | 4(52.28-53.65) | TFEC |
| XPEHH(HF); FST(Jersey) | 15(43.3-44.1) | TRIM66, STK33, DENND5A, SCUBE2, NRIP3 |
| XPEHH(Jersey); FST(BSW) | 16(21.1-22.9) | SPATA17, RRP15 |
| XPEHH(Jersey); iHS | 11(77.3-78.0) | TDRD15, APOB |
| XPEHH(Haryana);iHS | 3(64.1-65.8) | ADGRL4/ELTD1 |
| FST(BSW); iHS | 10(57.67-58.20) | MYO5A,bta-mir-1248-2, MYO5C |
| FST(HF); iHS | 14(24.36-24.62) | FAM110B,UBXN2B |
| FST(Haryana); iHS | 11(82.67-83.20) | DDX1, NBAS |

## 4. Discussion

The Admixture analysis and PCA plot reflects the presence of both indicine and taurine ancestry in our *Vrindavani* population, with a higher proportion of taurine ancestry (Holstein). The dominance of the Holstein component in *Vrindavani* cattle has also been recently reported in a different set of Vrindavani population (Ahmad et al., 2020).

In the present study, we wished to evaluate the effect of natural and artificial selection in *Vrindavani*, compared to its parent breeds. Due to genetic drift (Akey et al., 2002), and ascertainment bias of the SNP chip toward taurine breeds, it is difficult to distinguish true signatures of selection from false positives in crossbred cattle. Thus, three different methods of signature of selection (iHS, F_ST_ and XP-EHH) were applied with stringent thresholds to capture putative regions of selection across the genome.

The regions commonly detected by pairwise cross population methods (XP-EHH and F_ST_) against taurine breeds on BTA4 contain the *CACNA2D1* gene, which is a member of the calcium voltage-gated channel auxiliary subunit alpha-2 / delta, is previously reported to be a candidate gene associated with somatic cell score (Deng et al., 2011) and mastitis resistance (Yuan et al., 2010). Another gene on this chromosome is *TFEC*, reported to be a selection signature in African cattle and related with resistance to ticks and other tropical disease (Tijjani et al., 2019).

On BTA14, *FAM110B* and *UBXN2B* genes were identified to be associated with productive traits, reproductive traits (Grigoletto et al., 2019) and feed efficiency (Seabury et al., 2017). Flori et al. (2009) has also reported *FAM110* as selection signature for dairy cattle under artificial selection. The *STK33* gene on BTA15 reported in the F_ST_ and XP-EHH analyses against the Jersey and HF cattle, respectively, were reported as selection signatures in Gir cattle and are associated with milk production in indicine cattle (Maiorano et al., 2018).

Commonly identified regions from iHS and cross-population approaches against Hariana contain *TNNI3K* and *ADGRL4*/*ELTD1* genes on BTA 3. *TNNI3K* is a cardiac troponin interacting kinase, associated with udder depth (Kramer et al., 2014), inflammation mechanisms (Wiltshire et al., 2011) and lameness in Holstein–Friesian cattle (Sánchez-Molano et al., 2019). The *ADGRL4/ELTD1* gene is associated with milk fat yield (Li et al., 2010) and tick resistance (Porto Neto et al., 2011) in dairy cattle, average daily gain in pigs (Hong et al., 2017), and genetic predisposition for obesity in both humans and pigs (Lee et al., 2011). Another gene *DDX1* on BTA11 was reported to be involved in bovine mammary involution in environmental stress conditions (Dado-Senn et al., 2018). *DDX1* is also reported to be associated with linoleic acid content in Nellore cattle (Lemos et al., 2016) and viral resistance (Xue et al.,2019).

A common region in comparisons with both taurine and indicine breeds is detected on BTA2. It harbours *LRP1B* gene which codes for low density lipoprotein related with milk yield (Chen et al., 2105) and somatic cell score (Cole et al., 2011). *LRP* is widely expressed in several tissues and plays important roles in lipoprotein catabolism, blood coagulation, cell adhesion and migration (Haas et al., 2011).

## 5. Conclusion

This study provided the first overview of the selection footprints in the genome of the crossbred *Vrindavani* cattle of India. Results revealed that selection was operative more strongly in the regions related to environmental adaptation than milk yield, despite the latter being a focus of artifical selection. This could be explained by the presence of a large (50-75%) taurine inheritance in the Vrindavani genome, so a deviation from the parental breeds with respect to adaptation was not unexpected.

The slow rate of genetic gain with respect to the artificially selected productivity traits, due to the small and closed nature of the institutional herd examined in this study may also be responsible for our findings. Vrindavani is still a relatively new breed, and we expect these selection signatures to be more prominent in the coming generations. To further reduce false positives and increase the resolution of detection of selection signatures, we suggest validation of this study in a larger field herd using the HD genotyping array or whole-genome sequence data.

## Acknowledgements

The molecular work was supported by world bank funded CAAST-ACLH project of NAHEP.

## Data Availability Statement

The data that support the findings of this study are taken from open access public repository (http://widde.toulouse.inra.fr/widde/widde/main.do?module=cattle) (https://krishi.icar.gov.in/jspui/handle/123456789/31167) web portal. Additional data of *Vrindavani* cattle used in this study are available from the corresponding author on request.

## References

Ahmad, S.F., Panigrahi, M., Chhotaray, S., Pal, D., Parida, S., Bhushan, B., et al. (2020). Revelation of genomic breed composition in a crossbred cattle of India with the help of Bovine50K BeadChip. Genomics. 112, 1531–1535. doi: 10.1016/j.ygeno.2019.08.025

Akey, J.M., Zhang, G., Zhang, K., Jin, L. and Shriver, M.D. (2002). Interrogating a high-density SNP map for signatures of natural selection. Genome res. 12, 1805–1814. doi: 10.1101/gr.631202

Alexander, D.H., Novembre, J. and Lange, K. (2009). Fast model-based estimation of ancestry in unrelated individuals. Genome Res. 19, 1655–1664. doi: 10.1101/gr.094052.109

Amble, V.N. and Jain, J.P. (1967). Comparative performance of different grades of crossbred cows on military farms in India. J. Dairy Sci. 50, 1695–1702. doi: 10.3168/jds.S0022-0302(67)87696-3

Browning, B.L., Zhou, Y. and Browning, S.R. (2018). A one-penny imputed genome from next-generation reference panels. Am. J. Human Genet. 103, 338–348. doi: 10.1016/j.ajhg.2018.07.015

Chen, X., Cheng, Z., Zhang, S., Werling, D. and Wathes, D.C. (2015). Combining genome wide association studies and differential gene expression data analyses identifies candidate genes affecting mastitis caused by two different pathogens in the dairy cow. Open J. Anim. Sci. 5, 358–393. doi: 10.4236/ojas.2015.54040

Cheruiyot, E.K., Bett, R.C., Amimo, J.O., Zhang, Y., Mrode, R. and Mujibi, F.D. (2018). Signatures of selection in admixed dairy cattle in Tanzania. Front. Genet. 9, 607. doi: 10.3389/fgene.2018.00607

Chhotaray, S., Panigrahi, M., Pal, D., Ahmad, S.F., Bhanuprakash, V., Kumar, H., et al. (2019). Genome-wide estimation of inbreeding coefficient, effective population size and haplotype blocks in Vrindavani crossbred cattle strain of India. Biol. Rhythm Res. 1–14. doi: 10.1080/09291016.2019.1600266

Cole, J.B., Wiggans, G.R., Ma, L., Sonstegard, T.S., Lawlor, T.J., Crooker, B.A., et al. (2011). Genome-wide association analysis of thirty-one production, health, reproduction and body conformation traits in contemporary US Holstein cows. BMC genomics. 12, 408.

Dado-Senn, B., Skibiel, A.L., Fabris, T.F., Zhang, Y., Dahl, G.E., Peñagaricano, F. (2018). RNA-Seq reveals novel genes and pathways involved in bovine mammary involution during the dry period and under environmental heat stress. Sci. Rep. 8, 1–11. doi: 10.1038/s41598-018-29420-8

DAHD&F (2017-18). Department Of Animal Husbandry Dairying & Fisheries. Ministry Of Fisheries, Animal Husbandry & Dairying, New Delhi, India.

Danecek, P., Auton, A., Abecasis, G., Albers, C.A., Banks, E., DePristo, M.A., et al. (2011). The variant call format and VCFtools. Bioinformatics. 27, 2156–2158. doi: 10.1093/bioinformatics/btr330

Decker, J.E., McKay, S.D., Rolf, M.M., Kim, J., Alcalá, A.M., Sonstegard, T.S., et al. (2014). Worldwide patterns of ancestry, divergence, and admixture in domesticated cattle. PLoS Genet. 10, 1004254. doi: 10.1371/journal.pgen.1004254

Deng, G.X., Yuan, Z.R., Gao, X., Li, J.Y., Chen, J.B., Gao, H.J. et al. (2011). Identification mutation of the CACNA2D1 gene and its effect on somatic cell score in cattle. J. Appl. Anim. Res. 39, 15–18. doi: 10.1080/09712119.2011.558616

Flori, L., Fritz, S., Jaffrézic, F., Boussaha, M., Gut, I., Heath, S., et al. (2009). The genome response to artificial selection: a case study in dairy cattle. PloS one. 4, 6595. doi: 10.1371/journal.pone.0006595.

Gautier, M., Klassmann, A. and Vitalis, R. (2017). rehh 2.0: a reimplementation of the R package rehh to detect positive selection from haplotype structure. Mol. Ecol. Resour. 17, 78–90. doi: 10.1111/1755-0998.12634.

Grigoletto, L., Brito, L.F., Mattos, E.C., Eler, J.P., Bussiman, F.O., Silva, B.D.C.A., et al. (2019). Genome-wide associations and detection of candidate genes for direct and maternal genetic effects influencing growth traits in the Montana Tropical® Composite population. Livest. Sci. 229, 64–76. doi: 10.3389/fgene.2020.00123

Haas, J., Beer, A.G., Widschwendter, P., Oberdanner, J., Salzmann, K., Sarg, B., et al. (2011). LRP1b shows restricted expression in human tissues and binds to several extracellular ligands, including fibrinogen and apoE–carrying lipoproteins. Atherosclerosis, 216, 342–347. doi: 10.1016/j.atherosclerosis.2011.02.030

Hong, J.K., Jeong, Y.D., Cho, E.S., Choi, T.J., Kim, Y.M., Cho, K.H., et al. (2018). A genome-wide association study of social genetic effects in Landrace pigs. Asian Austral. J. Anim. 31, 784. doi: 10.5713/ajas.17.0440

Huang, D. W., Sherman, B. T., Zheng, X., Yang, J., Imamichi, T., Stephens, R., et al. (2009). Extracting biological meaning from large gene lists with DAVID. Curr. Protoc. Bioinformatics. 27, 13–11. doi: 10.1002/0471250953.bi1311s27

Kim, E.S. and Rothschild, M.F. (2014). Genomic adaptation of admixed dairy cattle in East Africa. Front. Genet. 5, 443. doi: 10.3389/fgene.2014.00443

Kramer, M., Erbe, M., Seefried, F.R., Gredler, B., Bapst, B., Bieber, A. et al. (2014). Accuracy of direct genomic values for functional traits in Brown Swiss cattle. J. Dairy Sci. 97, 1774–1781. doi: 10.3168/jds.2013-7054

Lee, K.T., Byun, M.J., Kang, K.S., Park, E.W., Lee, S.H., Cho, S., et al. (2011). Neuronal genes for subcutaneous fat thickness in human and pig are identified by local genomic sequencing and combined SNP association study. PloS one. 6, 16356. doi: 10.1371/journal.pone.0016356

Lemos, M.V., Chiaia, H.L.J., Berton, M.P., Feitosa, F.L., Aboujaoud, C., Camargo, G.M., et al. (2016). Genome-wide association between single nucleotide polymorphisms with beef fatty acid profile in Nellore cattle using the single step procedure. BMC genomics. 17, 213. doi: 10.1186/s12864-016-2511-y

Li, H., Wang, Z., Moore, S.S., Schenkel, F.S. and Stothard, P. (2010). Genome-wide scan for positional and functional candidate genes affecting milk production traits in Canadian Holstein Cattle. Proc 9th WCGALP, Leipzig, Germany, 26.

Maiorano, A.M., Lourenco, D.L., Tsuruta, S., Ospina, A.M.T., Stafuzza, N.B., Masuda, Y., et al. (2018). Assessing genetic architecture and signatures of selection of dual purpose Gir cattle populations using genomic information. PLoS One. 13, 0200694. doi: 10.1371/journal.pone.0200694

Porto Neto, L.R., Bunch, R.J., Harrison, B.E. and Barendse, W. (2011). DNA variation in the gene ELTD1 is associated with tick burden in cattle. Anim. Genet. 42, 50–55. doi: 10.1111/j.1365-2052.2010.02120.x.

Purcell, S., Neale, B., Todd-Brown, K., Thomas, L., Ferreira, M.A., Bender, D., et al. (2007). PLINK: a tool set for whole-genome association and population-based linkage analyses. Am.J. Hum. Genet. 81, 559–575. doi: 10.1086/519795

R Core Team (2018). R: A Language and Environment for Statistical Computing. Vienna: R Foundation for Statistical Computing.

Rocha, D., Billerey, C., Samson, F., Boichard, D. and Boussaha, M., 2014. Identification of the 366 putative ancestral allele of bovine single-nucleotide polymorphisms. J. Anim. Breed. Genet. 131, 483–486. doi: 10.1111/jbg.12095

Sabeti, P.C., Varilly, P., Fry, B., Lohmueller, J., Hostetter, E., Cotsapas, C., et al. (2007). Genome-wide detection and characterization of positive selection in human populations. Nature. 449, 913–918. doi: 10.1038/nature06250.

Sanchez-Molano, E., Bay, V., Smith, R.F., Oikonomou, G. and Banos, G., 2019. Quantitative trait loci mapping for lameness associated phenotypes in Holstein Friesian dairy cattle. Front. in Genet. 10, 926. doi: 10.3389/fgene.2019.00926

Seabury, C.M., Oldeschulte, D.L., Saatchi, M., Beever, J.E., Decker, J.E., Halley, Y.A., et al. (2017). Genome-wide association study for feed efficiency and growth traits in US beef cattle. BMC genomics. 18, 386. doi: 10.1186/s12864-017-3754-y

Singh, R.R., Dutt, T., Kumar, A., Tomar, A.K.S. and Singh, M. (2011). On-farm characterization of Vrindavani cattle in India. Indian J. Anim. Sci. 81, 267–271.

Tijjani, A., Utsunomiya, Y.T., Ezekwe, A.G., Nashiru, O. and Hanotte, O., 2019. Genome Sequence Analysis Reveals Selection Signatures in Endangered Trypanotolerant West African Muturu Cattle. Front. genet. 10, 442. doi: 10.3389/fgene.2019.00442

Voight, B.F., Kudaravalli, S., Wen, X. and Pritchard, J.K. (2006). A map of recent positive selection in the human genome. PLoS Biol. 4, 72. doi: 10.1371/journal.pbio.0040072

Weir, B.S. and Cockerham, C.C. (1984). Estimating F-statistics for the analysis of population structure. Evolution. 1358–1370. doi: 10.2307/2408641

Wiltshire, S.A., Leiva-Torres, G.A. and Vidal, S.M., 2011. Quantitative trait locus analysis, pathway analysis, and consomic mapping show genetic variants of Tnni3k, Fpgt, or H28 control susceptibility to viral myocarditis. J. Immunol. 186, 6398–6405. doi: 10.4049/jimmunol.1100159

Xue, Q., Liu, H., Zeng, Q., Zheng, H., Xue, Q. and Cai, X. (2019). The dead-box RNA helicase DDX1 interacts with the viral protein 3D and inhibits foot-and-mouth disease virus replication. Virolo. Sin. 34, 610–617. doi: 10.1007/s12250-019-00148-7

Yuan, Z.R., Li, J., Liu, L., Zhang, L.P., Zhang, L.M., Chen, C., et al. (2011). Single nucleotide polymorphism of CACNA2D1 gene and its association with milk somatic cell score in cattle. Mol. Boil. Rep. 38, 5179–5183. doi: 10.1007/s11033-010-0667-0

Zimin, A. V., Delcher, A. L., Florea, L., Kelley, D. R., Schatz, M. C., Puiu, D., et al. (2009). A whole-genome assembly of the domestic cow, Bos taurus. Genome Biol. 10, 42. doi:10.1186/gb-2009-10-4-r42.

